# Rigid Foot Soles Improve Balance in Beam Walking

**DOI:** 10.1101/510990

**Authors:** Meghan E. Huber, Enrico Chiovetto, Martin Giese, Dagmar Sternad

**Affiliations:** Department of Mechanical Engineering, Massachusetts Institute of Technology, Cambridge, Massachusetts, USA.; Section for Computational Sensomotorics, Department of Cognitive Neurology, Hertie Institute for Clinical Brain Research, Centre for Integrative Neuroscience, University Clinic Tübingen, Tübingen, Germany.; Departments of Biology, Electrical and Computer Engineering, and Physics, Northeastern University, Boston, Massachusetts, USA.

## Abstract

Maintaining balance while walking on a narrow beam is a challenging motor task. This is presumably because the foot’s ability to exert torque on the support surface is limited by the beam width. Still, the feet serve as a critical interface between the body and the external environment, and it is unclear how the mechanical properties of the feet affect balance. Here we examined how restricting the degrees of freedom of the feet influenced balance behavior during beam walking. We recorded whole-body joint kinematics of subjects with varying skill levels as they walked on a narrow beam with and without wearing flat, rigid soles on their feet. We computed changes in whole-body motion and angular momentum across these conditions. Results showed that wearing rigid soles improved balance in the beam walking task, but that practice with rigid soles did not affect or transfer to task performance with bare feet. The absence of any after-effect suggested that the improved balance from constraining the foot was the result of a mechanical effect rather than a change in neural strategy. Though wearing rigid soles can be used to assist balance, there appear to be limited training or rehabilitation benefits from wearing rigid soles.

## INTRODUCTION

Whether walking over rocks or across logs, humans have remarkable ability to maintain balance while navigating difficult terrain^1,2^. In fact, healthy humans are so proficient in their ability to balance that some turn to walking along a thin wire to truly challenge their skills. While there has been prolific research on the control of postural balance over the past decades, this work has largely focused on understanding how humans maintain balance during quiet standing^3–8^. Despite many insights into the limits of postural balance, it is still an open question how the central nervous system controls the highly redundant and extremely complex architecture of the body to maintain balance during more realistic locomotion, especially in challenging environments.

A paradox of human motor control is that while the human body is vastly complex (e.g., large number of degrees of freedom, long time delays, sensorimotor noise^9^, nonlinear muscle properties, intersegmental dynamics), the overt behavior is often surprisingly simple in structure. Thus, low-dimensional models, derived by compressing the number of degrees of freedom in the body, can be used to competently describe human balance. For example, an inverted pendulum can adequately capture much of the behavior that humans exhibit during quiet stance^8,10^. When the base of support is reduced, such as in the case of standing on a narrow beam, adding a second linkage to make a double-inverted pendulum model has proven sufficient^8,10,11^. In a recent study, Chiovetto et al.^12^ revealed that the segmental dynamics of the body, quantified by angular momentum of each segment about the beam, exhibited a lower-dimensional structure compared to the structure in relative joint angle kinematics. Using principal component analysis, a single component could explain about 90% of the variance in angular momentum of all body segments and was highly similar across individuals. Unlike several prior studies on standing and walking on a beam, Chiovetto et al.^12^ allowed participants to freely move their arms during the experiment with the goal to look at the full complexity of the realistic behavior. How does the nervous system control the high-dimensional architecture of the entire body to generate such low-dimensional patterns?

A critical aspect of maintaining balance is managing the physical interaction between the body and its external environment. Because the feet serve as interfaces through which the body and ground simultaneously act upon each other, they play a pivotal role in maintaining balance. As seen in the development of prosthetics, the mechanical properties of the foot can significantly influence balance behavior^13–15^. And yet, how the complex architecture of the human feet contributes to balance is still poorly understood. Each foot consists of many articulated, rigid segments which are surrounded by compliant, heterogeneous tissue, making it difficult to accurately measure and model the subtle coordinated behavior of the foot^16–18^. Paradoxically, most models of human balance drastically simplify the foot. In the inverted pendulum models of standing balance, the foot is typically reduced to a static, rigid segment attached to the ground acted upon by an ideal torque source at the ankle. Thus, the foot’s influence on human balance, particularly during walking, remains understudied.

The aim of this study was to understand how the degrees of freedom of the foot and the ankle contribute to maintaining mediolateral (ML) balance when walking on a narrow beam (Figure 1a). Both feet were constrained by attaching a flat, rigid sole to the bottom of each foot (Figure 1b). The rigid sole prevented any motion of the foot joints distal to the ankle, namely bending at the midfoot and torsion on the long axis of the foot. Importantly, plantarflexion/dorsiflexion and inversion/eversion ankle motion was not affected.

**Figure 1.**
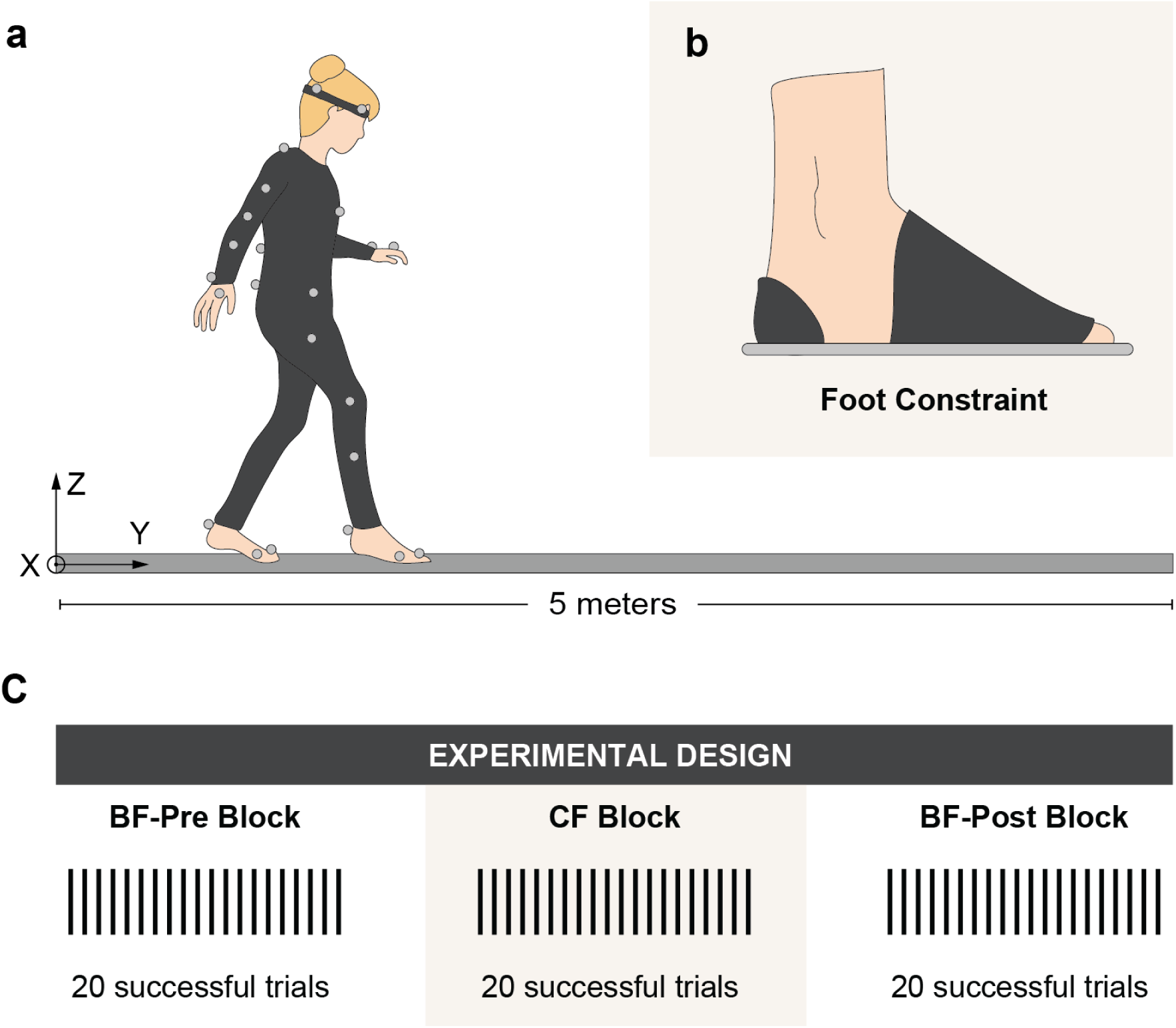
(a) In the experimental task, subjects walked on a narrow beam (3.4cm wide by 5m long) wearing reflective markers for 3D kinematic data acquisition. One traversal of the entire beam was considered a successful trial. (b) Flat, rigid soles were attached to the bottom of the subjects’ feet to constrain foot and toe bending. The dark regions indicate where soles were attached with hook and loop straps and reinforced with tape. Ankle motion was unaffected. (c) Subjects completed 20 successful trials in in each of the following three blocks: Bare Feet (BF)-Pre, Constrained Feet (CF), and Bare Feet (BF)-Post.

On the one hand, a highly flexible foot may be critical for actively sensing and controlling the physical interaction between the foot and the beam. Sawers et al.^19^ found that dancers have an increased set of available whole-body actions (i.e., more muscle synergies) to maintain balance when walking on a beam compared to novices. This underscores that limiting degrees of freedom reduces the number of movements available to withstand perturbations and maintain upright balance. Thus, constraining the set of motor actions of the feet could impair balance and worsen performance in the beam walking task (***Hypothesis 1a***). An alternative argument, however, is equally plausible. Constraining the foot to act as a rigid, flat segment could increase contact stability between the foot and the flat surface of the beam and thereby improve performance^20^. For example, Robbins et al.^21^ found that elderly men improved beam walking when they wore shoes with hard, thin soles. They stepped off the beam less frequently compared to performing the task with bare feet or shoes with softer soles. Hence, an alternative expectation is that rigid soles positively affect balancing performance (***Hypothesis 1b***).

Changing the mechanics of the feet could also cause subjects to adapt their control strategy for maintaining balance with practice. When the rigid soles are removed, this altered strategy could subsequently influence balance performance. For instance, if the rigid soles led to worse performance when the rigid soles were removed, we would expect subjects to quickly return to their original control strategy (***Hypothesis 2a***). This scenario corresponds many adaptation studies where, for example, the adaptation to a perturbing force field only persists as short-term after-effects as they are not functional when the perturbation is removed. If the adapted strategy leads to improved balance behavior after removing the soles, however, we would expect that this acquired strategy and its positive impact on performance would persist (***Hypothesis 2b***). This scenario would indicate that the soles acted as a teaching aid that could accelerate learning to balance. A third feasible scenario is that humans do not even alter their control policy when the rigid soles are attached to their feet. For example, if it is only the change in the foot mechanics that altered performance, we would not expect subjects to change their control policy (***Hypothesis 2c***). If this was the case, we would expect practice with constrained feet to have no influence on subsequent performance with bare feet. By assessing how practice of the beam-walking task with constrained feet influences subsequent balance behavior with bare feet, we gain insight not only into how the complex architecture of the foot influences the neural control of balance, but also whether this may be a suitable intervention for either assisting or rehabilitating impaired balance behavior.

This study investigated how constraining the foot affected mediolateral (ML) balance in beam walking for young individuals with varying levels of prior balance training. We tested whether constraining the feet influenced ML-balance during beam walking compared to performing the task with bare feet. Previous work has shown that the velocity of the center of mass (COM-V) in the ML-direction is a good indicator of skilled balance^12^. Hence, impaired balance is indicated by an increased velocity of the center of mass (COM-V) in the ML-direction and increased whole-body angular momentum (WB-AM) about the beam axis; improved balance would show the opposite trend. To evaluate whether practice with constrained feet affected performance after removing the rigid soles, we tested subjects walking with bare feet before and after walking with rigid soles. In addition to testing the hypotheses, further analyses of whole-body coordination were conducted to shed light on *how* constraining the foot influenced ML-balance during beam walking.

The results showed that constraining the feet improved ML-balance in the beam walking task (***Hypothesis 1b**)*. Moreover, task performance with bare feet was unaffected by practice with rigid soles (***Hypothesis 2c***). Together, these findings indicate that the improvement in balance from constraining the foot was the result of a mechanical effect rather than a change in neural strategy. Additional analyses showed that the angular momentum of most individual segments was reduced when wearing the rigid soles. Moreover, the contribution of ankle torque relative to hip torque was increased when the feet were constrained. We propose that constraining the feet improved performance because of an increase in contact stability between the foot and the beam^20^.

## RESULTS

Seven healthy subjects took part in the experiment. Their prior balance training ranged from none to several years in competitive gymnastics. In each trial, subjects were instructed to walk the length of a narrow beam (3.4cm wide and 5m long) without stepping off the beam (Figure 1a). A trial was deemed successful if the subject did not step off before reaching the end of the beam; otherwise the trial was declared a failed trial. Subjects had to complete 20 successful trials in each of the following three blocks: The first block consisted of 20 successful trials with bare feet (BF-Pre block), followed by 20 successful trials with constrained feet (CF block), and another 20 successful trials with bare feet (BF-Post block) (Figure 1c).

### Number of Failed Trials

To gauge if constraining subjects’ feet affected their ability to accomplish the beam-walking task, we examined its influence on the number of failed trials in each of the three blocks. A one-way within-subject analysis of variance (ANOVA) revealed that foot condition (BF-Pre, CF, BF-Post) did not have a significant effect on the number of failed trials (F_2,12_ = 0.38, p = 0.69) (Figure 2). On average, subjects failed in approximately 4-5 trials in each block. As expected, performance across subjects varied along a continuum determined in part by their prior balance training. Subjects who exhibited the best performance (shown in red and orange in Figure 2) were trained gymnasts. As the results below show, the cohort presented a sufficient spectrum of balance abilities that allowed more general conclusions.

**Figure 2.**
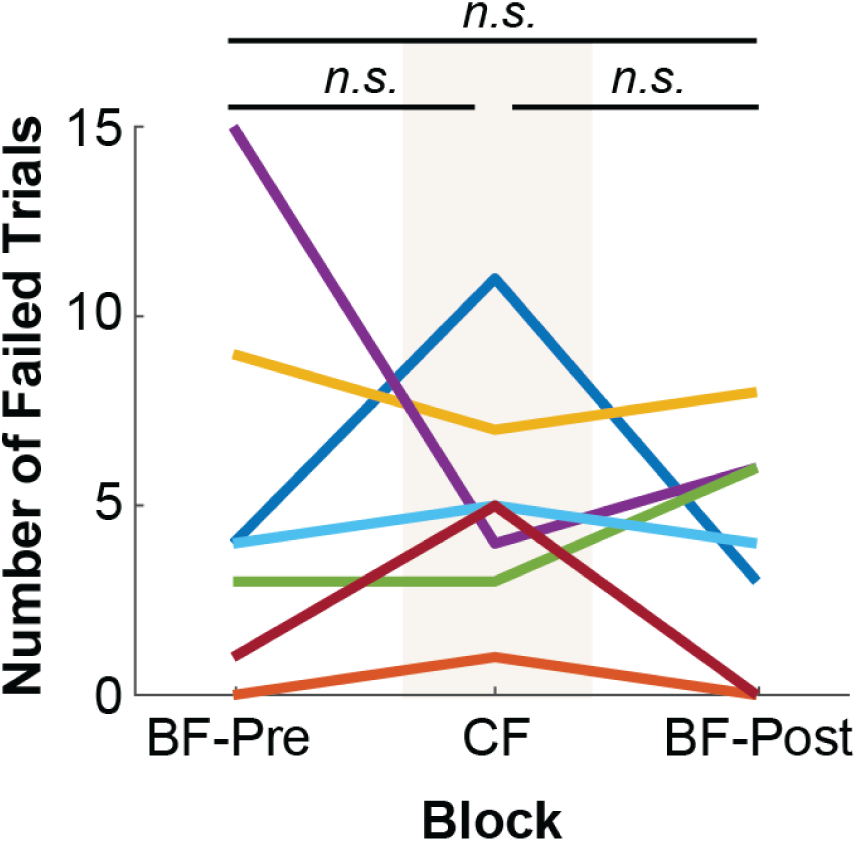
Number of failed trials in each block of individual subjects. “*n.s.”* indicates no significant difference. The orange and red lines with the overall lowest number of failed trials indicate the two gymnasts.

### Example Data

Even though constraining the subjects’ feet did not require more attempts to accomplish the overall task goal, analysis of more fine-grained measures revealed that it did significantly influence their balance proficiency as they performed the task. Figure 3a-c displays the series of body postures of two representative subjects during a typical trial in each of the three conditions. For reference, data from Example Subject 1 is shown in light blue in other results figures; Example Subject 2, who was trained in gymnastics, is shown in dark red. Subjects displayed not only large trunk movements, but also large and variable movements of both arms. Importantly, these body movements were visibly reduced in the CF block. Figure 4a-b shows the time series of the center of mass velocity (COM-V) and whole-body angular momentum (WB-AM) for the corresponding trials shown in Figure 3a-c. Performance in each trial was summarized by taking the root-mean-square (RMS) of these dependent measures in the middle 67% (i.e., two-thirds) of the trial. For both subjects, the amplitude of the COM-V and WB-AM signals were decreased in the CF block (Figure 4a-b), reflecting the decrease in body movement observed in Figure 3b.

**Figure 3.**
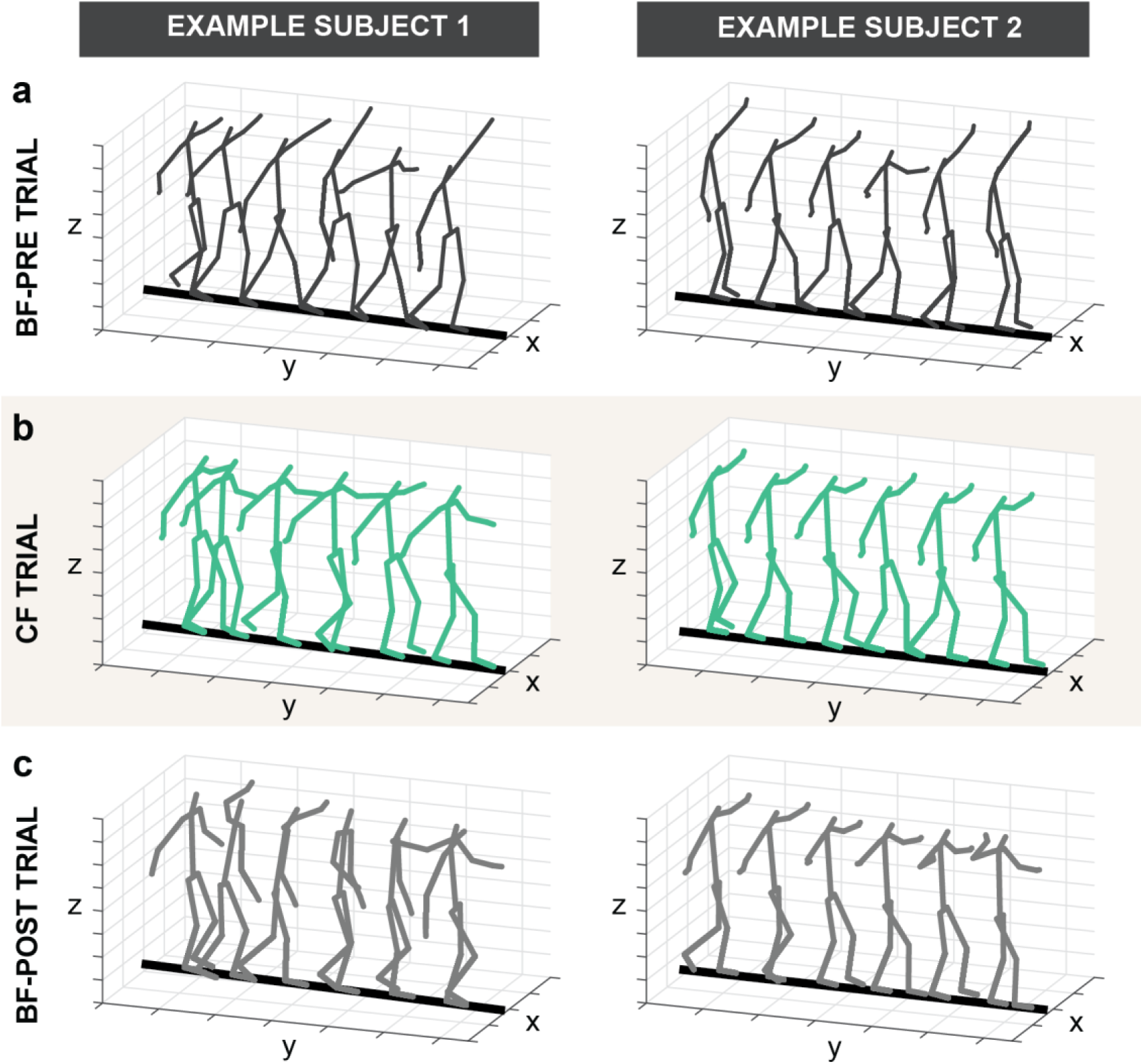
Representative stick figures of two example subjects walking across the beam constructed from 3D kinematic data during representative trials in each block: (a) BF-Pre, (b) CF, and (c) BF-Post. Subject 1 pertains to the blue lines and subject 2 belongs to the red line in Figure 2.

**Figure 4.**
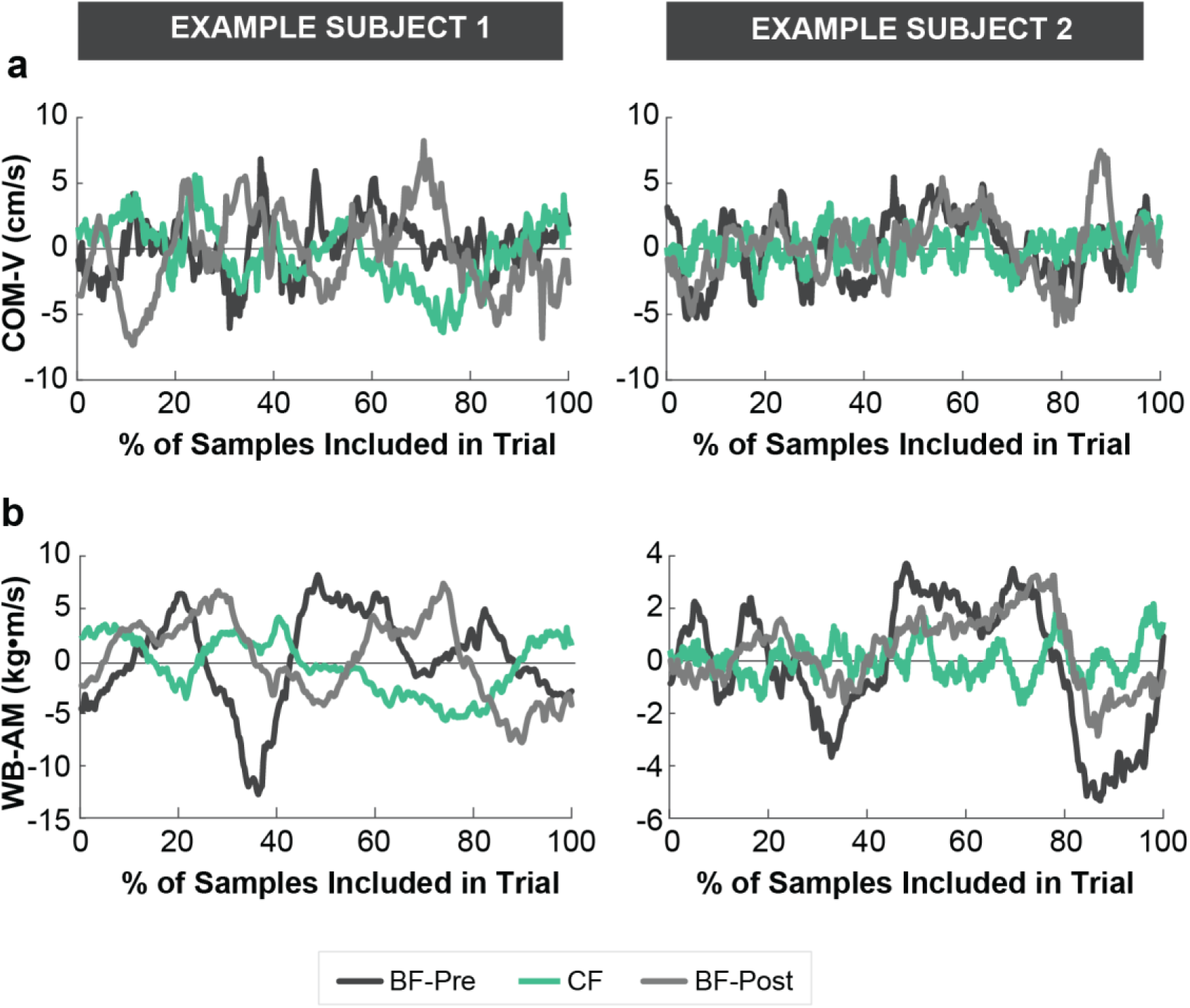
Time series of (a) COM-V and (b) WB-AM of representative trials in each block from the two example subjects. The trials shown here are the same as in Figure 3. Only the middle 67% of samples (indicated in this figure as 100% of included samples) were used for calculating the root-mean-square (RMS) of each dependent measure in each trial.

### Center of Mass Velocity (COM-V)

As demonstrated in prior work^12^, the RMS of COM-V in the ML direction of each trial is sensitive to differences in balance proficiency. Consistent with qualitative observations from Figure 4a, a one-way within-subject ANOVA revealed a significant effect of block on RMS of COM-V (F_2,12_ = 32.99, p = 0.000031) (Figure 5a-b). Planned comparisons showed a significant reduction in RMS of COM-V from the BF-Pre block (M = 2.93cm/s, SD = 0.68cm/s) to the CF block (M = 2.09cm/s, SD = 0.55cm/s) (t_6_ = 8.40; p = 0.00016). This was followed by a significant increase from the CF block to the BF-Post block (M = 2.92cm/s, SD = 0.74cm/s) (t_6_ = −6.93; p = 0.00045) (Figure 5b). These results indicated that constraining subjects’ feet significantly improved their ML-balance (***Hypothesis 1b***). Moreover, there was no difference in the mean RMS of COM-V between the BF-Pre block and the BF-Post block (t_6_ = 0.06, p = 0.95) (Figure 5b). Even though subjects varied in their ability to perform the task, the decrease in the RMS of COM-V during the CF block was seen across all subjects.

**Figure 5.**
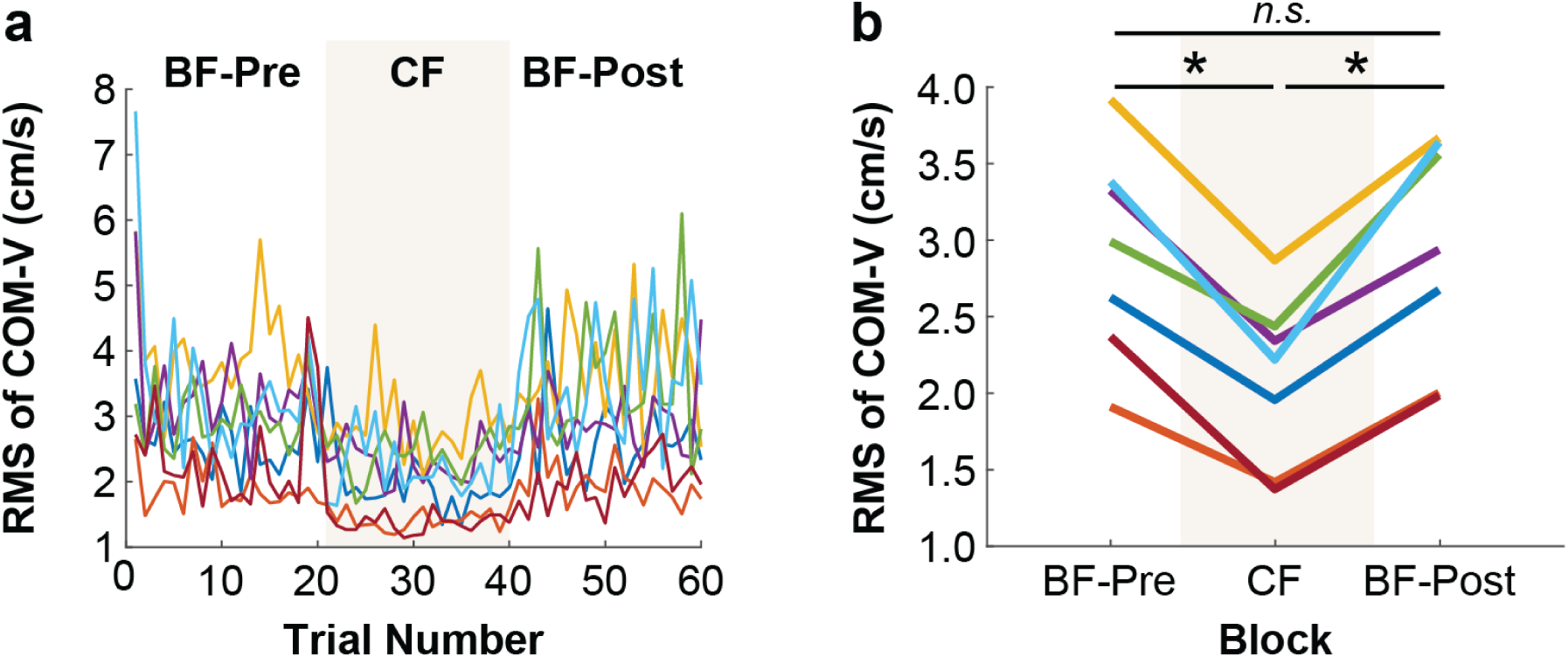
RMS of COM-V of individual subjects (a) across all trials and (b) averaged within each block. An asterisk represents p < 0.05, and “*n.s.”* indicates no significant difference.

To address Hypothesis 2a-c, the last successful trial of the BF-Pre block was compared with the first trial of BF-Post block. There was no significant difference between the last trial of the BF-Pre block (M = 2.80cm/s, SD = 0.66cm/s) and the first successful trial of the BF-Post block (M = 2.62cm/s, SD = 0.71cm/s) (t_6_ = 0.52, p = 0.62), indicating an immediate return to baseline performance after the rigid soles were removed (Figure 5a). Thus, practice with constrained feet did not influence subjects’ balance performance with bare feet (***Hypothesis 2c***).

### Whole-Body Angular Momentum (WB-AM)

We also examined how performing the balance beam task with rigid soles influenced subjects’ whole-body angular momentum (WB-AM) about the axis of the beam. The measure of WB-AM quantified the angular momentum of a subject’s body with respect to the beam. In the beam walking task, the body was subject to ground reaction forces acting on the feet. These external forces induced considerable changes in the body’s WB-AM. We quantified WB-AM with respect to the beam axis, rather than the body’s center of mass or head position for two reasons: First, the beam was fixed and thus provided an inertial reference frame. Second, our prior work revealed that the structure of AM was less complex when quantified about the beam axis^12^.

The same one-way ANOVA rendered a significant effect of block on the RMS of WB-AM (F_2,12_ = 21.73, p < 0.001) (Figure 6a-b). Planned comparisons revealed that constraining the foot had a similar effect on RMS of WB-AM as it did on COM-V. The RMS of WB-AM significantly decreased from the BF-Pre block (M = 5.07kg·m^2^/s, SD = 2.40kg·m^2^/s) to the CF block (M = 3.17kg·m^2^/s, SD = 1.52kg·m^2^/s) (t_6_ = 4.69, p = 0.0034), and then significantly increased from the CF block to the BF-Post block (M = 4.75kg·m^2^/s, SD = 2.21kg·m^2^/s) (t_6_ = −5.33, p = 0.0018) (Figure 6b). There was no difference in RMS of WB-AM between the BF-Pre block and the BF-Post block (t_6_ = 1.71, p = 0.14) (Figure 6b), nor between the last successful trial of the BF-Pre block (M = 4.01kg·m^2^/s, SD = 2.08kg·m^2^/s) and the first successful trial of the BF-Post block (M = 4.67kg·m^2^/s, SD = 2.158kg·m^2^/s) (t_6_ = −0.86, p = 0.42), (Figure 6a). Again, these results indicate that constraining subjects’ feet significantly improved their ML balance (***Hypothesis 1b***), but the improved performance with constrained feet did not transfer or influence subjects’ subsequent performance with bare feet (***Hypothesis 1c***).

**Figure 6.**
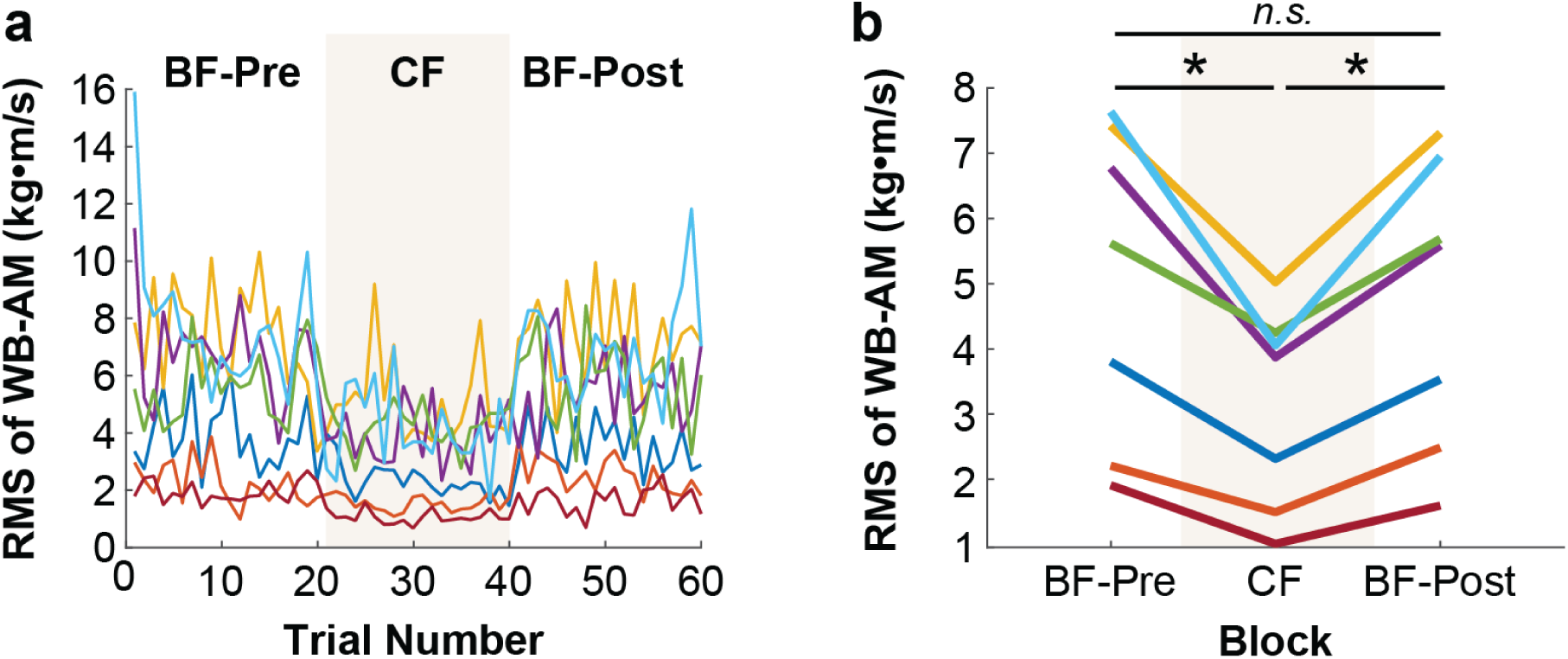
RMS of WB-AM of individual subjects (a) across all trials and (b) averaged within each block. An asterisk represents p < 0.05, and “*n.s.”* indicates no significant difference.

To further understand how WB-AM was reduced, we also examined how angular momentum (AM) of the individual body segments changed when wearing rigid soles. As WB-AM was calculated as the sum of AM from individual body segments about the beam, it could have been lowered in several different ways. For example, WB-AM could have been decreased by reducing the AM of either some or all body segments, by changing the direction of segments’ AM relative to each other, or even a combination of both. Thus, we next assessed how wearing rigid soles influenced the spatiotemporal patterns of the individual body segments in their contribution to WB-AM.

### Angular Momentum (AM) of Individual Body Segments

For each of the 15 body segments, a one-way within-subject ANOVA was conducted on the RMS of each segment’s AM. The results of each ANOVA are detailed in Table 1. To summarize, the effect of block on each segment’s AM was significant, except for the left and right feet. When significant, planned comparisons revealed that the effect of block on each segment’s AM was similar to its effect on WB-AM. Each segment’s AM significantly decreased from the BF-Pre block to the CF block (ps > 0.014) and then subsequently increased from the CF block to the BF-Post block (ps < 0.024) (Figures 7-8). There were no significant differences between AM in the BF-Pre and BF-Post blocks (ps > 0.14). Hence, the reduction in WB-AM when wearing rigid soles was due in large part to a reduction in each segment’s contribution to WB-AM. It was not the result of reduced AM from a single large segment, for example.

**Table 1.** One-way within subject ANOVA results on individual body segment AM with block as independent variable

| Segment # | Dependent Measure | dF | SS | F | p |
| --- | --- | --- | --- | --- | --- |
| 1 | Pelvis AM | 2 | 0.89 | 22.39 | 0.000089* |
| 2 | Right Thigh AM | 2 | 0.32 | 22.85 | 0.000081* |
| 3 | Left Thigh AM | 2 | 0.31 | 18.30 | 0.00023* |
| 4 | Right Shank AM | 2 | 0.16 | 9.69 | 0.0031* |
| 5 | Left Shank AM | 2 | 0.021 | 9.93 | 0.0029* |
| 6 | Right Foot AM | 2 | 0.00053 | 0.87 | 0.45 |
| 7 | Left Foot AM | 2 | 0.00054 | 2.16 | 0.16 |
| 8 | Head AM | 2 | 2.96 | 20.22 | 0.00014* |
| 9 | Thorax/Ab AM | 2 | 6.13 | 10.92 | 0.0020* |
| 10 | Right Upper Arm AM | 2 | 0.31 | 12.35 | 0.0012* |
| 11 | Left Upper Arm AM | 2 | 0.41 | 8.71 | 0.0046* |
| 12 | Right Forearm AM | 2 | 0.43 | 10.60 | 0.0022* |
| 13 | Left Forearm AM | 2 | 0.58 | 9.29 | 0.0037* |
| 14 | Right Hand AM | 2 | 0.23 | 11.49 | 0.0016* |
| 15 | Left Hand AM | 2 | 0.25 | 10.16 | 0.0026* |

\*statistically significant

**Figure 7.**
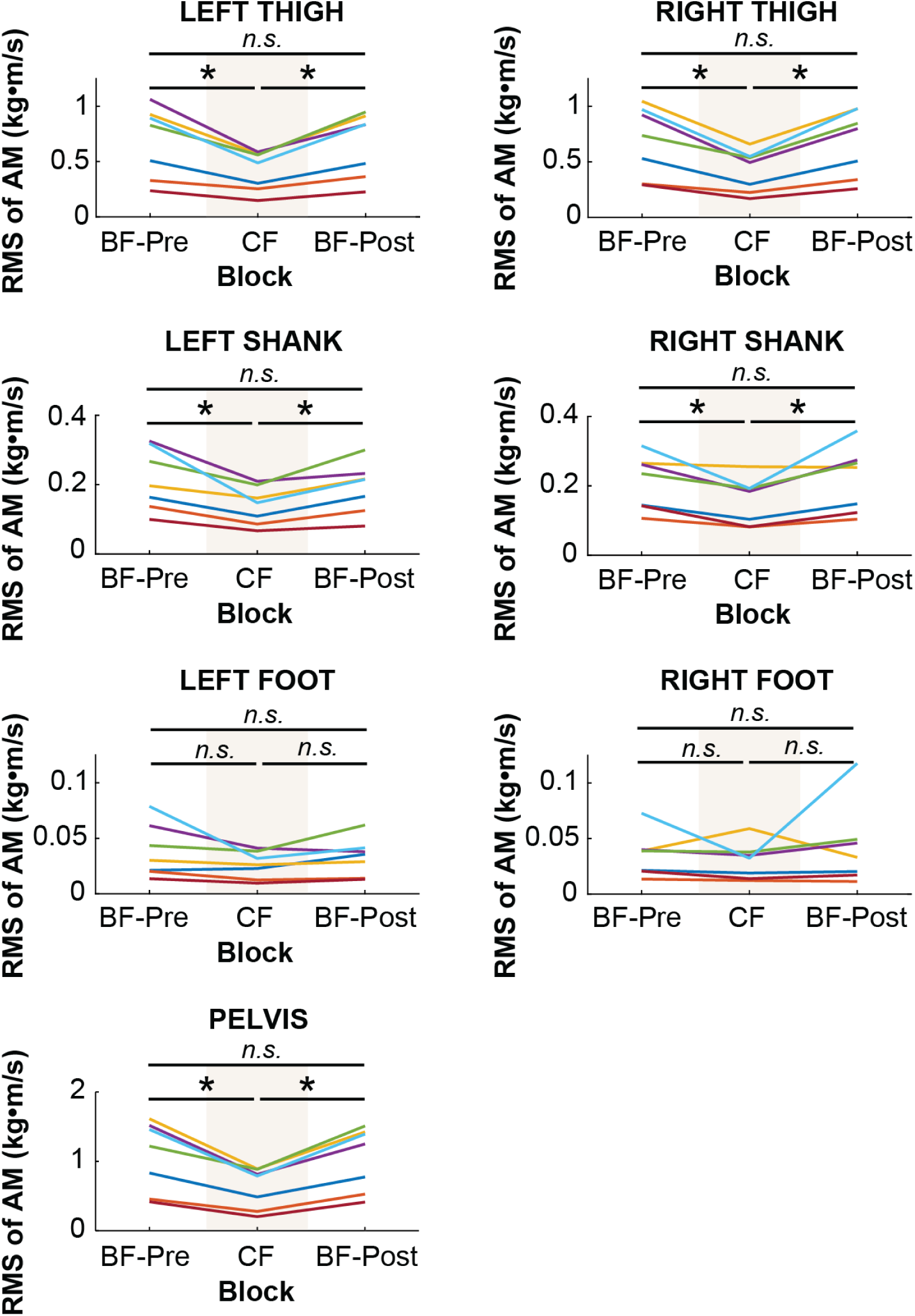
RMS of lower body segment AM of individual subjects averaged within each block. An asterisk represents p < 0.05, and “*n.s.”* indicates no significant difference.

**Figure 8.**
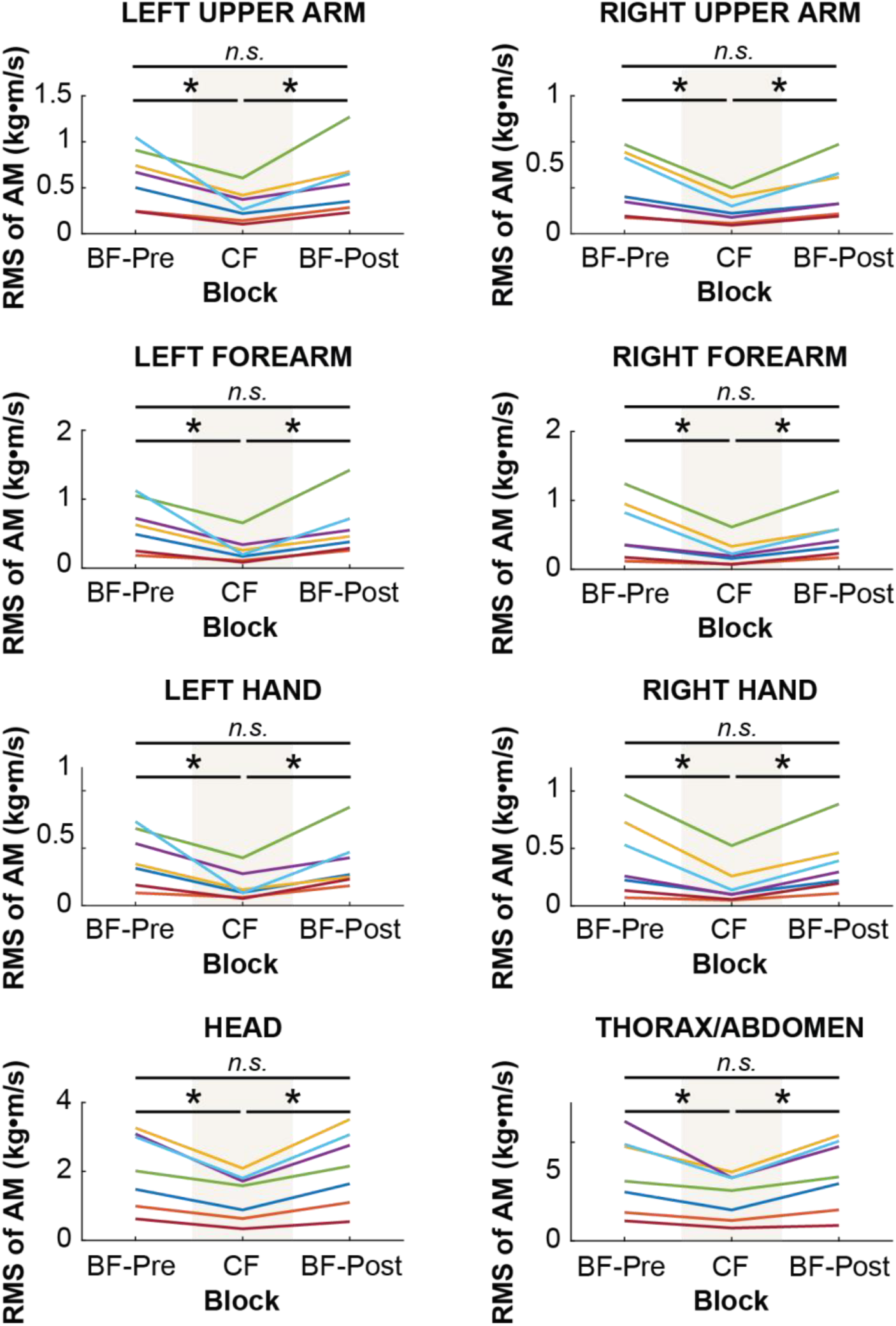
RMS of upper body segment AM of individual subjects averaged within each block. An asterisk represents p < 0.05, and “*n.s.”* indicates no significant difference.

### Correlation of Upper-and Lower-Body Angular Momentum (CORR-AM)

As seen in Figure 9 the upper-body segments (head, thorax, upper arms, lower arms, and hands) generated AM opposite in direction to the AM generated by the lower-body segments (pelvis, thighs, shanks, and feet). To examine if the coordination of upper-body and lower-body AM contributions were affected by constraining the foot, we computed correlation between the sum of AM of lower body segments (LB-AM) and the sum of AM of upper-body segments (UB-AM) for each trial, which we refer to as CORR-AM. Note that three outlier trials (0.7% of all trials) were omitted from the analysis as the CORR-AM values were uncharacteristically low. Consistent with the representative data shown in Figure 9, upper-body AM and lower-body AM were highly anti-correlated as the overall mean of CORR-AM across all conditions was −0.88 (SD = 0.05). As illustrated in Figure 10, rotation of the upper body segments (with respect to the hip) was opposite to that of the lower body segments (with respect to the beam). This suggests that subjects used a “hip-dominant” strategy to maintain balance.^22^

**Figure 9.**
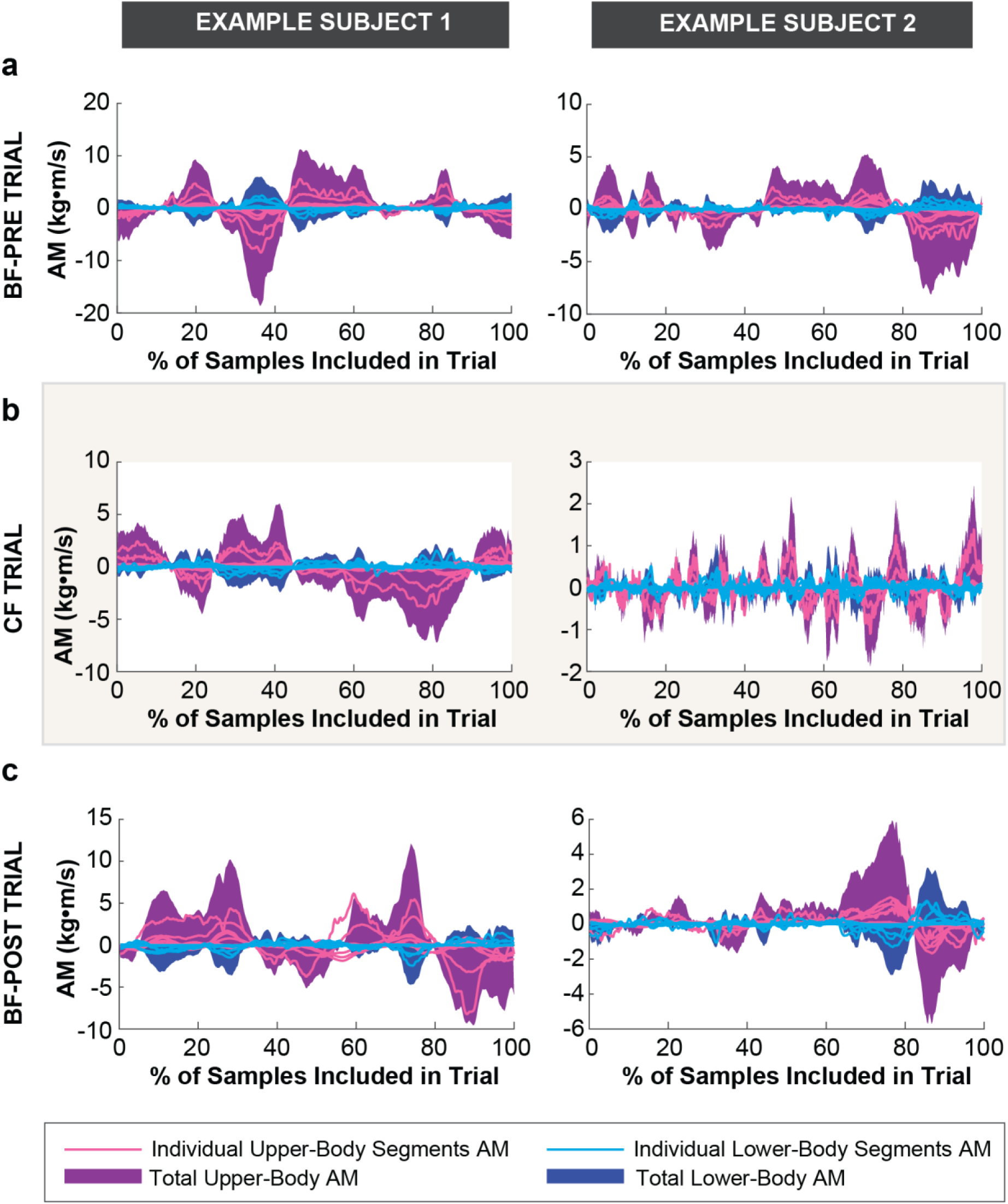
Time series of individual body segment AM, total UB-AM, and total LB-AM of representative trials from two example subjects in each block: (a) BF-Pre, (b) CF, (c) BF-Post. Only the middle 67% of samples (indicated as 100% of included samples in this figure) were used for calculating the root-mean-square (RMS) of individual segment AM measure and correlation between UB-AM and LB-AM (CORR-AM) in each trial.

**Figure 10.**
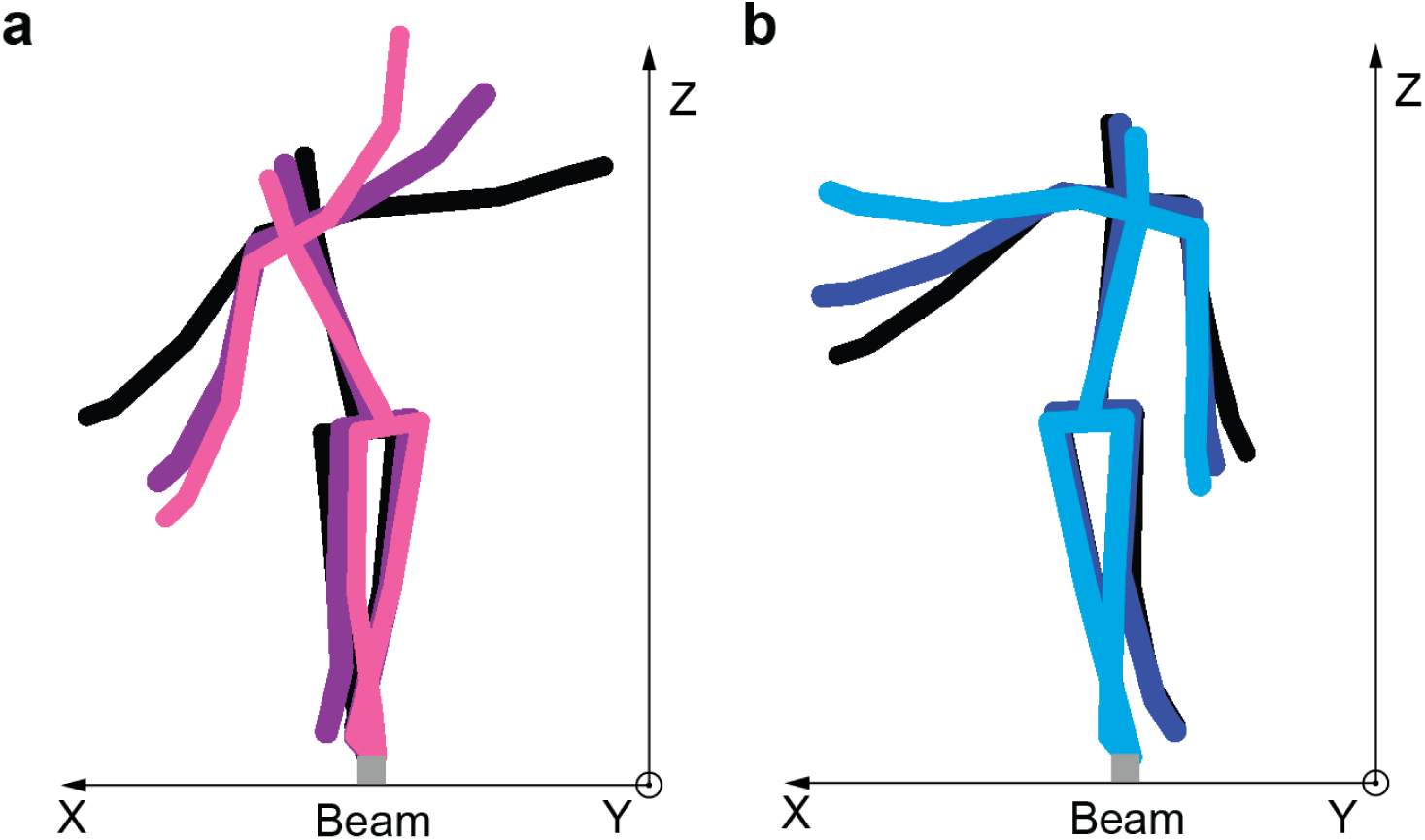
Representative stick figures of Example Subject 1 walking along the beam shown in the frontal plane. The color of the skeletons becomes lighter as the subject progresses forward along the beam axis (i.e., y-axis). (a) When the lower body segments rotated clockwise about the beam, the upper segments rotated counterclockwise about the hip. (a) When the lower body segments rotated counterclockwise about the beam, the upper segments rotated clockwise about the hip.

A one-way within-subjects ANOVA found a significant effect of block on CORR-AM (F_2,12_ = 22.75, p = 0.000083) (Figure 11a-b). The CORR-AM significantly increased, i.e., became less correlated, from the BF-Pre block (M = −0.89, SD = 0.03) to the CF block (M = −0.84, SD = 0.06) (t_6_ = −4.89, p = 0.0027) and then significantly decreased, i.e., became more correlated, from the CF block to the BF-Post block (M = −0.90, SD = 0.04) (t_6_ = 5.45, p = 0.0016) (Figure 11b). There was no difference in CORR-AM between the BF-Pre block and the BF-Post block (t_6_ = 0.85, p = 0.43) (Figure 11b), nor between the last successful trial of the BF-Pre block (M = −0.88, SD = 0.06) and the first successful trial of the BF-Post block (M = −0.94, SD = 0.03) (t_6_ = 1.84, p = 0.12), (Figure 11a). Interestingly, the upper-body AM and lower-body AM became less correlated when the foot was constrained, even though balance proficiency was improved. In fact, the trained gymnasts (red and orange traces in Figure 11a-b) also had the least anti-correlation.

**Figure 11.**
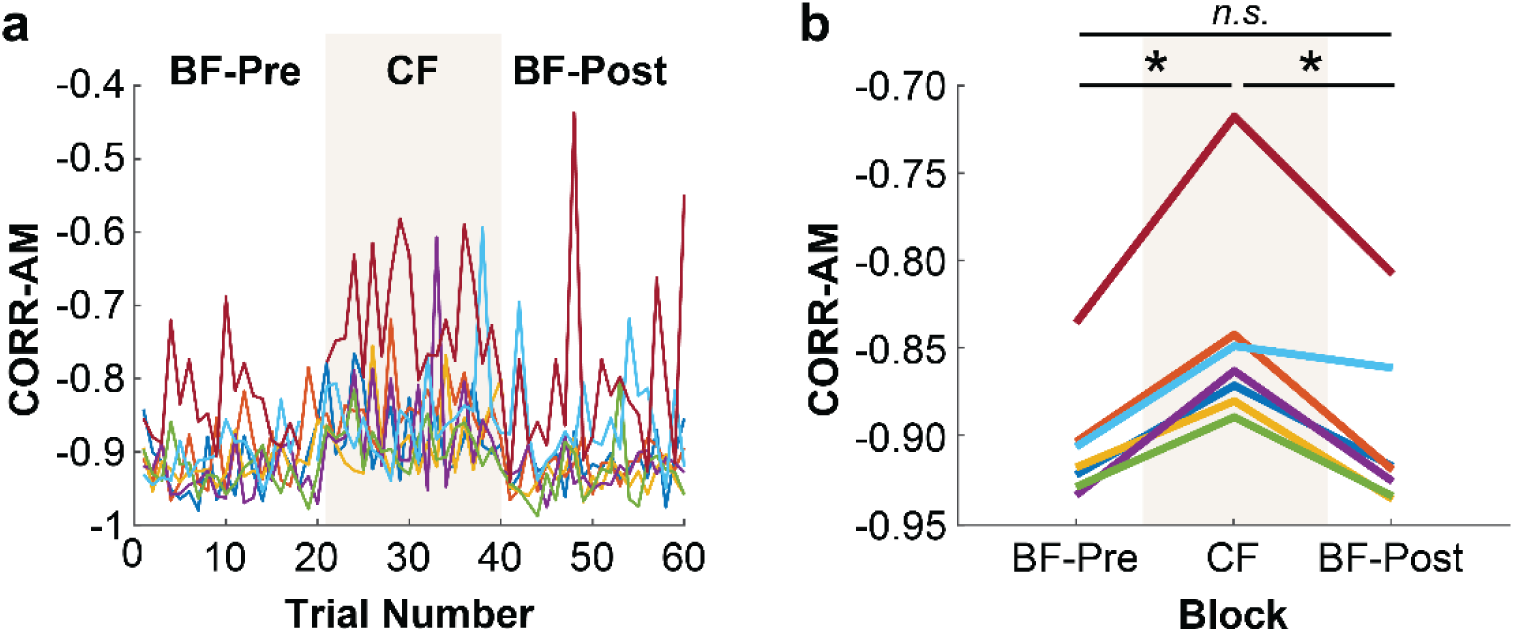
CORR-AM of individual subjects (a) across all trials and (b) averaged within each block. An asterisk represents p < 0.05, and “*n.s.”* indicates no significant difference.

## DISCUSSION

The goal of this study was to determine how reducing the complexity of the foot influenced whole-body coordination to maintain balance in a challenging beam walking task. We found that constraining the feet with rigid soles immediately improved balance as indicated by a reduction in the variability of COM-V and WB-AM. However, we did not find evidence that subjects altered their control strategy in response to the reduction in foot degrees of freedom. Practicing the task with rigid soles did not influence subsequent behavior with bare feet.

In support of **Hypothesis 1b** our results showed that constraining the feet with rigid soles is an effective method of assisting balance. This finding is in accordance with the conclusions of Robbins et al.^21^. By further assessing the effect of practice with the rigid soles, however, we found that it may not be an effective intervention for training or rehabilitating balance (**Hypothesis 2c**). In general, an intervention meant to enhance the performance or learning of a motor skill and should change neural control such that it results in improved task performance under normal conditions^23,24^. In our study, barefoot performance was unaffected by practice with constrained feet. In fact, we did not even observe a short-lived after-effect, which further suggests that subjects did not change their neural control strategy. While it is conceivable that subjects could alter their control strategy with long-term practice wearing rigid soles^19^, it remains an open question whether that learned strategy would improve or impair subsequent barefoot performance. It also cannot be ruled out that subjects might have learned a new control strategy, but that strategy was entirely context-dependent such that it did not transfer^25^. Evidence for this would require testing the effect of longer practice with the rigid soles to seek improvements within one condition. Without any further investigations, the results presented here suggest that constraining the feet may not be an effective intervention for training or retraining balance. Understanding how this manipulation improved balance performance can inform the development of future interventions.

When balancing on a narrow base of support, the ankle’s ability to exert torque on the beam through the foot is limited^10^. Thus, it is not surprising that numerous studies have reported that humans use a hip-dominant strategy to maintain balance when standing on a beam^3,4,8,10,22,26^. Consistent with these studies of standing balance, we similarly observed that subjects used a hip strategy to maintain ML balance when walking on a beam, as indicated by high anti-correlation between the AM generated by the upper-and lower-body segments^22^. Importantly, this was observed even though the arms were allowed to move freely, representative of real-world conditions. Even though subjects used a hip-dominant strategy during beam walking, this did not mean that the influence of the foot and ankle was minimal as is often presumed when balancing on a narrow beam. In fact, our finding that constraining the feet significantly altered balance behavior showed otherwise.

Not only did constraining the feet decrease the overall AM magnitude of most individual segments, it also resulted in less anti-correlation between the AM of the upper-and lower-body segments. Though the change in anti-correlation was small, it was significant and observed in all subjects (Figure 11b). As demonstrated in a prior simulation study of a double-inverted pendulum model^22^, the degree of anti-correlation decreases when the overall magnitude of ankle torque increases relative to the magnitude of hip torque. These simulations assumed no change in signal-to-noise ratio. It is important to note that the increase in relative ankle contribution observed with constrained feet could have resulted from increased ankle torque, decreased hip torque, or a combination of the two. While we cannot definitively discern how the change in relative ankle contribution occurred, we do know that it can be attributed to altering the physical interaction between the foot and beam. The fact that subjects did not appear to learn or adopt a new control strategy during practice with constrained feet further supports the notion that the improvement in balance was the result of a mechanical effect.

We speculate that constraining the feet improved balance because it increased the stability of contact between the foot and the beam. Note that the width of the support surface was identical in the BF and CF conditions, meaning that adding the flat, rigid soles did not increase the maximum torque that could be applied at the ankle. However, constraining the feet may have increased the “effective” range of ankle torque. While the multiple degrees of freedom in each foot may increase control and/or sensing abilities, they also make the foot compliant. Without the soles, exerting large ankle torque onto the beam could cause the compliant foot to rotate. If so, subjects possibly reduced the amount of torque applied at the ankle to avoid this rotation. Future studies comparing the distribution of pressure under the feet in each condition would shed further light on this possible explanation.

Wearing rigid soles may have increased the amount of torque that could be applied at the ankle without resulting in foot rotation about the beam^20^, and thus improved performance. But note, this is only one mechanism through which the “effective” range of ankle torque could have been increased. Interestingly, the subjects who were most proficient at maintaining balance tended to have less anti-correlation of upper and lower body momentum in the BF conditions. It is possible that these subjects were either able to modulate the mechanical properties of their ankle and foot or they could better compensate for the interaction dynamics at the foot-beam contact. This could explain why Sawers and Ting^19^ observed more muscle synergies in experienced balancers. For instance, reducing the interaction dynamics could require finer control of the degrees of freedom in the feet (e.g., toes) that expert dancers and balancers might learn with training. This also underscores that simple structure in overt balance behavior is not necessarily indicative of a “simple” controller in the neuromotor system.

While our results gave clear evidence that adding flat rigid soles can assist balance, this benefit to balance may come at cost. For instance, Takahashi et al.^27^ found that wearing shoes with stiff soles significantly increased the metabolic cost of walking. Moreover, we observed that there was no transfer from practicing beam walking with constrained feet to walking with bare feet. Ultimately, future work is needed to further understand (1) the influence of the foot and ankle mechanical properties on balance, and (2) how expert balancers modulate or compensate for its effects. We expect that addressing these open questions will yield promising new insights for enhancing the assistance and rehabilitation of balance.

## METHODS

### Subjects

Seven healthy subjects (gender: 2 females and 5 males, age: 28.7 ± 2.5 years, mass: 68.4 ± 10.9 kg, height: 1.74 ± 0.08 m) took part in the experiment. None had any prior experience with the specific experimental task. The experiment conformed to the Declaration of Helsinki and written informed consent was obtained from all participants according to the protocol approved by the ethical committee at the Medical Department of the Eberhard-Karls-Universität of Tübingen, Germany.

### Experimental Protocol

In each trial, the subject walked along a narrow beam (3.4 cm wide, 3.4 cm high, 4.75 m long) at a self-selected speed. Before the start of each trial, subjects stood with their left foot on the beam and their right foot on the ground. After the experimenter gave the “go”-signal, they placed their right foot on the beam and began walking. Upon reaching the end of the beam, subjects were instructed to step off, placing their feet on either side of the beam. Subjects did not receive any other instruction on how to walk or how fast they should walk across the beam. They could use all body segments, including arms, as they wished to maintain balance. For data processing, the placement of the right foot on beam indicated the start of each trial; the last step before stepping off the beam marked the end of the trial. A trial was deemed successful, if the subject remained on the beam for its entire length. If the subject lost balance and had to step on the ground before reaching the end, the trial was labeled as unsuccessful. After each trial, subjects were allowed to take a short rest if needed.

Each subject was instructed to complete 20 successful trials in each of the following three blocks: Bare Feet–Pre (BF–Pre), Constrained Feet (CF), and Bare Feet–Post (BF–Post). In the BF–Pre and BF–Post blocks, participants walked without shoes; in the CF block, participants performed trials with flat, rigid soles attached to each foot. The solid soles were 3D printed and designed to be slightly larger than all subjects’ feet (width: 12cm at widest point, length: 31cm, depth: 1cm). All subject wore the same size soles. They were secured to the subjects’ feet with hook and loop straps and reinforced with duct tape as illustrated in Figure 1b. These soles did not affect the plantar/dorsi-flexion and inversion/eversion motion of the ankle.

### 3D Motion Capture Data Collection

Reflective markers were placed on the subjects’ bodies following Vicon’s Plug-In Gait marker set (Figure 1). During each trial, 3D whole-body motion capture data was collected using a 10-camera motion capture system (Vicon, Oxford, UK) at a sampling rate of 100Hz. As illustrated in Figure 1a, the origin of the lab coordinate frame was set to the start end of the beam, with its y-axis aligned along the beam and its x-axis perpendicular to the beam. Commercial Vicon software was used to reconstruct and label the markers to interpolate between short missing segments in the 3D marker trajectories.

Based on the subject’s self-reported height and weight, subject-specific dynamic models (Plug-In Gait model consisting of 15 rigid body segments, Table 1) were fit to the 3D marker trajectories using C-Motion Visual3D software (Germantown, MD). The dependent measures for each trial were calculated using the model-based data exported from Visual3D that were subsequently analyzed using custom scripts in Matlab (The Mathworks, Natick, MA) as described in detail below.

### Dependent Measures

#### Number of Failed Trials

The number of failed trials for each subject in each block presented a course-grained measure for overall task performance.

#### Center of Mass Velocity (COM-V)

For each sample in a given trial, the COM-V in the ML-direction (i.e., *x*-direction) was approximated by computing the backward difference of the COM position values. The COM position in the ML-direction (i.e., along the x-axis) at sample *t*, denoted as *r*_*x*_(*t*), was calculated as follows:

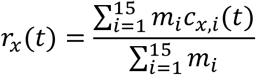

where *m*_*i*_ denoted the mass of the *i*^th^ segment, *c*_*x,i*_(*t*) was the position of the *i*^th^ segment’s COM in the *x*-direction at sample*t*, and *n* was the total number of samples in each trial. For each trial, the RMS of COM-V was calculated using COM-V values only samples from the middle 67% (i.e., two-thirds) of each trial. This was also done for all subsequent RMS calculations to avoid any possible transients or fatigue effects in the dependent measures. For each subject, the average RMS COM-V across trials in each block was calculated to assess change in balance performance across blocks.

#### Individual Body Segment Angular Momentum (AM)

In each trial, the angular momentum (AM) from each of the 15 body segments was calculated individually. In a given trial, the AM of *i*-th body segment at each sample *t* about the beam axis (i.e., *y*-axis), denoted by *L*_*y,i*_(*t*), was calculated using the following equation:

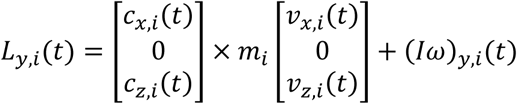

where *c*_*x,i*_(*t*) and *c*_*z,i*_(*t*) are the positions and *v*_*x,i*_(*t*) and *v*_*z,i*_(*t*) are the linear velocities of the *i*-th segment’s center of mass in the respective *x*-and *z*-directions at sample *t*. (*Iω*)_*y,i*_(*t*) was the angular momentum of the *i*-th segment about its center of mass in the lab coordinate frame. Note that AM of segment *i* about its center of mass was first calculated in its local coordinate frame and then transformed into the lab coordinate frame using the Visual3D software. For each trial, the RMS of the AM from each body segment was then calculated, again only for samples from the middle 67% of each trial.

#### Whole-Body Angular Momentum (WB-AM)

WB-AM about the beam axis (i.e., *y*-axis) at sample *t* in a given trial, denoted by *L*_*y,wb*_(*t*), was calculated by summing as the AM of all individual body segments as follows:

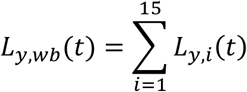

For each trial, the RMS of the AM from each body segment was then calculated using only samples from the middle 67% of each trial.

#### Correlation of Upper-and Lower-Body Angular Momentum (CORR-AM)

The correlation coefficient at lag-0 was calculated between the lower-body and upper-body AM signals in each trial. Lower-and upper-body AM about the beam axis (i.e., *y*-axis) at sample *t* in a given trial, denoted respectively by *L*_*y,lb*_(*t*) and *L*_*y,ub*_(*t*), were calculated by summing as the AM of the lower-and upper-body segments, respectively, as follows:

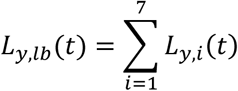

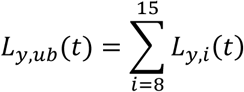

Only the middle 67% of samples in the signals of each trial were used to calculate the correlation coefficient.

### Statistical Analyses

To assess change in balance performance and strategy across blocks, the average RMS of COM-V, RMS of WB-AM, RMS of AM from individual body segments, and CORR-AM across trials in each block was calculated for each subject. These measures, along with the number of failed trials, were then subject to a one-way within-subjects ANOVA with block (BF-Pre, CF, BF-Post) as the within-subject factor. Planned comparisons in the form of pairwise t-tests were conducted to further assess the effect of block, when significant. The assumption of sphericity in the observed dependent measures was not violated as indicated by the non-significant results from Mauchly’s tests in all ANOVAs conducted.

To determine whether practice in the CF block had an immediate influence on subsequent balance performance with bare feet, a pairwise t-test was conducted to compare RMS of COM-V, RMS of WB-AM, and CORR-AM in the last trial of the BF-Post block and the first trial of the BF-Post block average RMS of COM-V.

In all statistical tests, the significance level was set to p < 0.05. The ANOVAs were performed using SPSS Statistics for Windows, Version 24.0 (IBM Corporation, Armonk, NY), and the pairwise t-tests were performed using MATLAB, Version 2016b (The Mathworks, Natick, MA).

## ACKNOWLEDGEMENTS

MEH was supported by the Max Planck Institute for Intelligent Systems, Tübingen. EC and MG were supported by BW Stiftung NEU007/1, BMBF CRNC FKZ 01GQ1704, EC H2020 ICT-23-2014 /644727 CogIMon, HFSP RGP0036/2016, DFG GZ: KA 1258/15-1. DS was supported by NSF-CRCNS-1723998, NSF-NRI-1637854, NIH-R01-HD087089 and the Max Planck Institute for Intelligent Systems, Tübingen.

We thank Prof. Neville Hogan, Prof. Stefan Schaal, Prof. Ludovic Righetti, Dr. Alexander Herzog, and Jongwoo Lee for their fruitful discussions and valuable feedback. We also thank Winfried Ilg for his help with the motion capture system.

## AUTHOR CONTRIBUTIONS

All authors conceived and designed the experiments. E.C. and M.E.H. carried out the experiments and processed the data. M.E.H. analyzed the data and drafted the manuscript. All authors discussed the results and implications. M.E.H. and D.S. edited the manuscript. All authors completed and approved the final version of the manuscript.

## DATA AVAILABILITY

The data will be made available on the website of Dagmar Sternad: https://web.northeastern.edu/actionlab/research/.

